# FAS: Assessing the similarity between proteins using multi-layered feature architectures

**DOI:** 10.1101/2022.09.01.506207

**Authors:** Julian Dosch, Holger Bergmann, Vinh Tran, Ingo Ebersberger

## Abstract

**Motivation:** Expert curation to differentiate between functionally diverged homologs and those that may still share a similar function routinely relies on the visual interpretation of domain architecture changes. However, the size of contemporary data sets integrating homologs from hundreds to thousands of species calls for alternate solutions. Scoring schemes to evaluate domain architecture similarities can help to automatize this procedure, in principle. But existing schemes are often too simplistic in the similarity assessment, many require an a-priori resolution of overlapping domain annotations, and those that allow overlaps to extend the set of annotations sources cannot account for redundant annotations. As a consequence, the gap between the automated similarity scoring and the similarity assessment based on visual architecture comparison is still too wide to make the integration of both approaches meaningful.

**Results:** Here, we present FAS, a scoring system for the comparison of multi-layered feature architectures integrating information from a broad spectrum of annotation sources. Feature architectures are represented as directed acyclic graphs, and redundancies are resolved in the course of comparison using a score maximization algorithm. A benchmark using more than 10,000 human-yeast ortholog pairs reveals that FAS consistently outperforms existing scoring schemes. Using three examples, we show how automated architecture similarity assessments can be routinely applied in the benchmarking of orthology assignment software, in the identification of functionally diverged orthologs, and in the identification of entries in protein collections that most likely stem from a faulty gene prediction.

**Availability and implementation:** FAS is available as python package: https://pypi.org/project/greedyFAS/

## 1 Introduction

The sequencing of genomes from organisms representing even the remotest corners of the tree of life is in full swing (Lewin et al., 2018; Mukherjee et al., 2021; Sayers et al., 2021). A rich toolbox exists to integrate the proteins encoded in these genomes into a comprehensive evolutionary network of significantly similar sequences (e.g. (Altschul et al., 1997; Buchfink et al., 2014; Potter et al., 2018; Steinegger & Söding, 2017)), and to differentiate between homologs originating from gene duplications (paralogs) and from speciation events (orthologs) (e.g. (Altenhoff et al., 2019; Sonnhammer & Östlund, 2015)). As orthologs tend to overlap at least partly in their function, this provides at the same time a tentative functional annotation for the newly identified genes/proteins (Fang et al., 2010; Gabaldón & Koonin, 2013). To facilitate a better-informed functional annotation *in silico*, dedicated databases compile, annotate and model evolutionary conserved protein domains, e.g., Pfam (Mistry et al., 2021), SMART (Letunic et al., 2021), HAMAP (Pedruzzi et al., 2015), CDD (Lu et al., 2020), or InterPro (Blum et al., 2021). They further provide the infrastructure to scan proteins for their domain content and order. The resulting linear arrangement of domains from N- to C-terminus, the domain architecture (DA), is informative about protein function (Bashton & Chothia, 2007; Burge et al., 2012; Doğan et al., 2016; Forslund & Sonnhammer, 2008; Kummerfeld & Teichmann, 2009; Messih et al., 2012), and many databases provide next to orthology assignments also the domain architectures as an accessory information (*e.g*., Sonnhammer & Östlund, 2015; Zdobnov et al., 2021). A visual interpretation of these DAs can indicate evolutionary changes with a likely effect on protein function (e.g. (Birikmen et al., 2021; Hsu et al., 2016; Huang et al., 2012; Gerrard & Bornberg-Bauer, 2003; Moore et al., 2014)). In turn, tools like Blast2GO (Conesa & Götz, 2008), BlastKOALA/GhostKOALA (Kanehisa et al., 2016) or eggNOG-mapper (Cantalapiedra et al., 2021) exploit automated DA similarity assessments to increase the specificity of functional annotation transfers between homologs for thousands of proteins.

DA similarity measures that allow a large-scale tracing of domain architecture changes between homologous proteins followed by manual curation of interesting candidates are provided by various scoring schemes. The Jaccard index, *i.e*., the intersection of domains annotated in two proteins over their union, is the simplest among these measures (Geer et al., 2002). More elaborated solutions consider additionally the extent of domain order conservation in the compared architectures using either the conservation of neighboring domain pairs or a DA alignment. They assess the similarity in position of shared domains, the agreement in copy number for individual domains, and they optionally weigh the contribution of individual domains to the overall similarity score (Doğan et al., 2016; Koestler et al., 2010; Lee & Lee, 2009; K. Lin et al., 2006; Song et al., 2007). Most of these approaches analyse linear DAs where each amino acid residue is assigned to at most one protein domain. Ambiguous assignments that arise, for example, from a sequence displaying a significant similarity to more than one Pfam model are typically resolved by selecting the domain with the lowest e-value (Lewis et al., 2019; Yeats et al., 2010). Koestler et al. devised the first scoring scheme that naturally handles overlapping domain annotations. Moreover, it allowed to extend DAs to multi-layered feature architectures (MLFA) where each layer represents information from a different annotation source (Koestler et al., 2010).

The comparison of MLFAs reduces the risk that two architectures with the same overlapping domain annotations are considered dissimilar due to reciprocal domain selection during overlap resolution (Koestler et al., 2010). However, the abstention of resolving overlapping domains can also result in underestimating architecture similarities, which generates a spurious signal of functional diversification (Fig. 1). Thus, there is still no satisfying solution for the scoring of pairwise feature architecture similarities.

**Figure 1:**
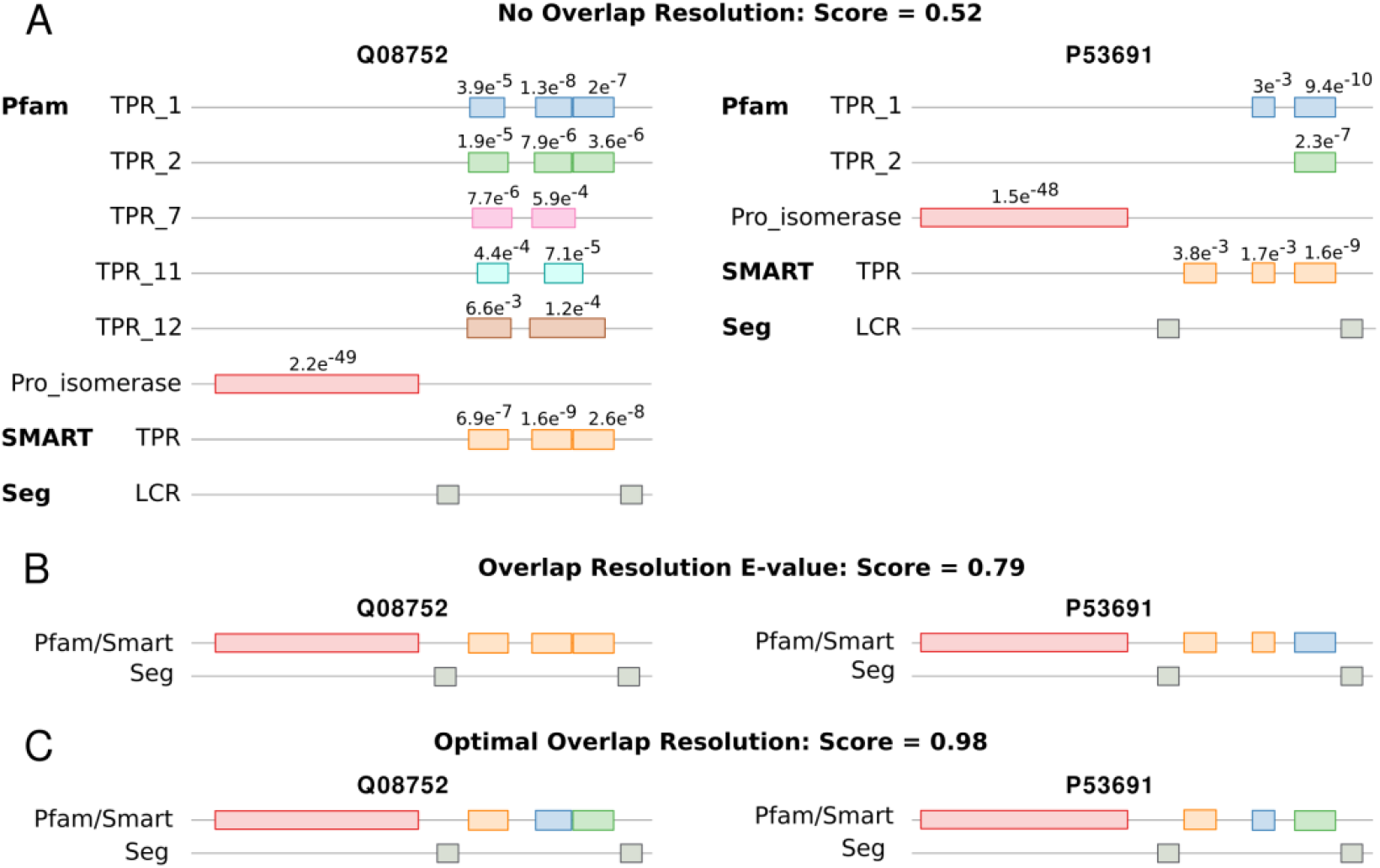
Multi-layered feature architectures of the human-yeast orthologous protein pair Q08752 and P53691. Both proteins are annotated as a Peptidyl-prolyl cis-trans isomerase D. Where available, e-values of the feature annotations are given above the respective feature instances. (A) Assessing the pairwise similarities of the two partly redundant feature architectures results in a similarity score of only 0.52. (B) Resolving the redundant feature architectures using a minimal e-value criterion results in a score of 0.79. (C) The optimal overlap resolution achieves a score of 0.98

Here, we introduce *FAS* to assess the similarities of MLFAs between protein pairs. As a main innovation, FAS resolves overlapping, redundant annotations by identifying the highest scoring linear path within one annotation layer. We show that this results in consistently higher similarity scores when compared to an e-value-informed overlap resolution. At the same time, it avoids scoring biases arising from the comparison of unresolved architectures causing both under- and overestimation of similarity scores. We demonstrate the use of the FAS scoring for benchmarking orthology assignment tools. We show that low FAS scores indicate lower semantic similarities of the GO annotation between the compared proteins, and we propose the detection of spurious gene annotations in newly sequenced genomes as a new use case for feature architecture similarity assessments. In summary, FAS substantially narrows the gap between automatically generated similarity scores and the assessment based on visual inspection. It therefore paves the way to routinely integrate feature architecture similarity scoring into the assessment of functional similarity between evolutionarily related proteins.

## 2 Concept and Implementation

### 2.1 Terminology

A *feature* represents a substring of protein *S* that can be assigned to one *feature class*. We differentiate distinct feature classes, where each class comprises features from one annotation source (see section 2.2). The union of all feature classes constitutes the *feature space*. Each feature class comprises one to many *feature types*. A Pfam family represented by its profile Hidden Markov model (pHMM) is one example of a feature type from the class “Pfam”. Each feature type is then represented by zero to many *instances*. The union of all annotated feature instances ordered along the protein sequence resembles the *feature architecture* of a protein *S* where each feature class corresponds to one *layer* in the architecture. *Overlaps* in the feature architecture are generated if a sequence position is assigned to two or more types of the same feature class. We consider a feature architecture as (partly) redundant if at least one overlap extends to *k* or more successive amino acids. A representative feature architecture is the realization of a non-redundant architecture by overlap resolution that maximizes the pairwise similarity score.

### 2.2 Feature space

The default feature space of FAS comprises Pfam and SMART domains (Letunic et al., 2021; Mistry et al., 2021), transmembrane domains annotated with tmhmm (Krogh et al., 2001), low complexity regions (Harrison, 2017) and coiled coils, predicted with COILS2 (Lupas, 1996). Alternatively, FAS can use architectures resulting from an InterPro scan annotation (Blum et al., 2021). The feature space can be adjusted by adding/removing features classes, and details are provided in the software manual.

### 2.3 Scoring feature architecture similarity

The feature architecture similarity score (FAS score) has been initially introduced by Koestler et al. 2010 and will briefly recapitulated in the following. It is a non-symmetric measure of the feature architecture similarity between two proteins, the reference *S* and the target *O*. The score ranges from a minimum of 0, when the two architectures have no feature type in common, to a maximum of 1, when the reference architecture resembles a (sub-)architecture of the target. The FAS score is a linear combination of (i) the multiplicity score (MS) capturing the fraction of feature types in the architecture of the reference protein that is represented in the target protein, and (ii) the positional score (PS) that captures the similarity of the position for the shared features in the two proteins. We compute

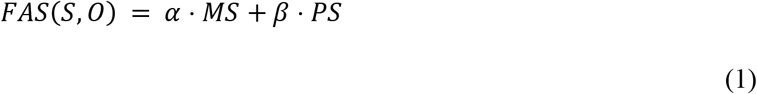

We use a default parameterization of *α=0. 7* and *β=0.3* to give the shared presence of features in two domain architectures a higher impact on the FAS score than their relative position in the proteins. However, we allow for case-by-case adaptation of these weights by the user.

#### 2.3.1 Multiplicity Score

We define *N* as the non-redundant set of feature types annotated in *S*, where each feature type *N_i_* occurs with 1 to *m* instances in the architecture of *S*. We compute the multiplicity score as

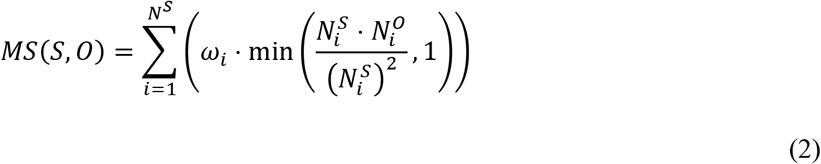

In brief, for each *N_i_* we multiply the number of instances of that feature in the reference 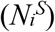 and the ortholog 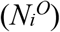 and normalize the product by the square number of feature instances in the reference. If 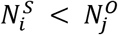, we apply an upper bound of 1. The contribution of each feature to the MS is then weighted by a factor *ω_i_* (see 2.3.3) such that the maximum of MS is 1.

#### 2.3.2 Positional Score

Let 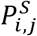 the relative position of the *jth* instance of feature type *N_i_* in *S*. We then identify the corresponding instance *l* of *N_i_* in *O*, such that the relative distance of the two instances in *S* and *O* is minimized. We compute the positional score as

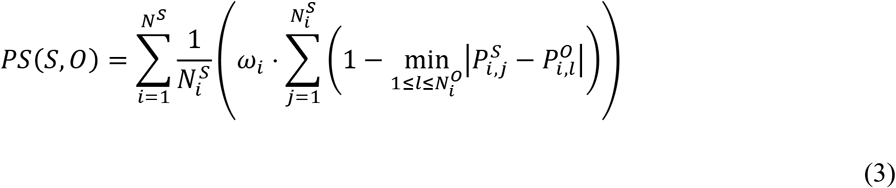

The relative positions 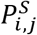 and 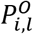 are computed as the absolute position of the feature instance midpoint in the protein sequence divided by the length of the protein. We sum the weighted partial scores for all features in *N* such that the PS reaches a maximum of 1.

#### 2.3.3 Domain weighting

The influence of individual feature types to the MS and the PS, respectively, is controlled by two main weighting schemes. In the uniform weighting scheme, all feature types contribute equally to the scoring. In the non-uniform weighting, we increase the weight of a feature *i* with its decreasing abundance in the proteome of the reference species. Per default, we calculate the weight of a feature as

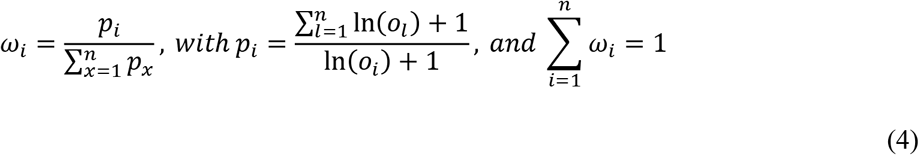

where *o_i_* is the number of instances of feature *i* in the reference proteome, and *n* is the sum of all feature types in the currently scored path. In contrast to Koestler et al. 2010, we compute the natural logarithm of the feature abundance for the calculation of *p* rather than using the raw feature abundance. This results in a less steep drop of weights for features with a higher abundance and increases their contribution to the score (Fig. S1). Next to the natural logarithm, users can choose from three alternative functions (log10, root4, root8) to further modulate the abundance-driven weighting (see Fig. S1). We further provide the option to specify a lower bound for the weights of individual feature types.

### 2.4 Resolution of redundant architectures

The main innovation of FAS is the resolution of redundant parts in the feature architecture. We represent the multi-layered feature architecture of a protein as a directed, acyclic graph (Fig. 2). Vertices denote the annotated feature instances, and edges connect the vertices such that the order in the graph reflects their order in the protein sequence. Per default, we connect only features of the same feature class, with one exception. Both SMART and Pfam domains are represented by profile hidden Markov models (pHMMs), and many SMART domains overlap with corresponding Pfam domains. Consequently, we subsume both in one feature class. First, we simplify the graph by ignoring overlaps of length smaller than *k* (we use a default of *k*=10). A path through the graph originates at the *start*, visits vertices with increasing distance from the start and terminates in the *end* vertex. Redundant annotations are resolved by selecting one of the alternative feature types. From all alternative paths through redundant parts of a feature architecture, we then aim to identify the one that maximizes the pairwise FAS score. We have implemented two alternative search strategies.

**Figure 2:**
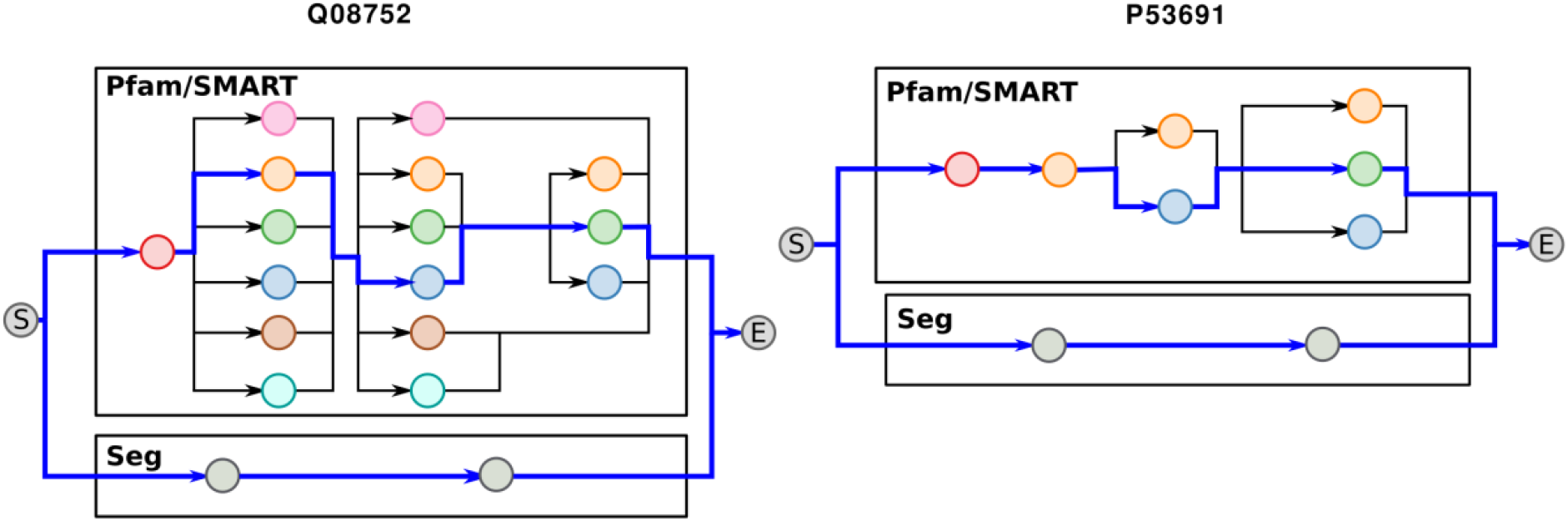
Representation of the feature architectures of Q08752 and P53691 as directed acyclic graphs. Each vertex represents a feature instance shown in Fig. 1 using the same coloring. The architecture comprises two annotation layers, Pfam/SMART, and SEG. Branching within a layer indicates redundant annotations. The blue path identifies the highest scoring linear path through the Pfam/SMART layer

#### 2.4.1 Exhaustive path search

FAS traverses the graph in an exhaustive depth-first search scoring each path of the target protein against each alternative path of the reference protein in the same annotation layer. The best scoring pair of paths is then added to the representative architectures. In the example shown in Fig 3, there are 72 alternative paths through the architecture of the human protein, and only 6 paths through that of the yeast protein. Evaluating all possible 72×6=432 alternative combinations reveals that the optimal resolution of the redundant parts in the feature architecture results in a FAS score of 0.98 compared to only 0.79 that was achieved with a naïve resolution using an e-value cutoff (see Fig. 1).

**Figure 3:**
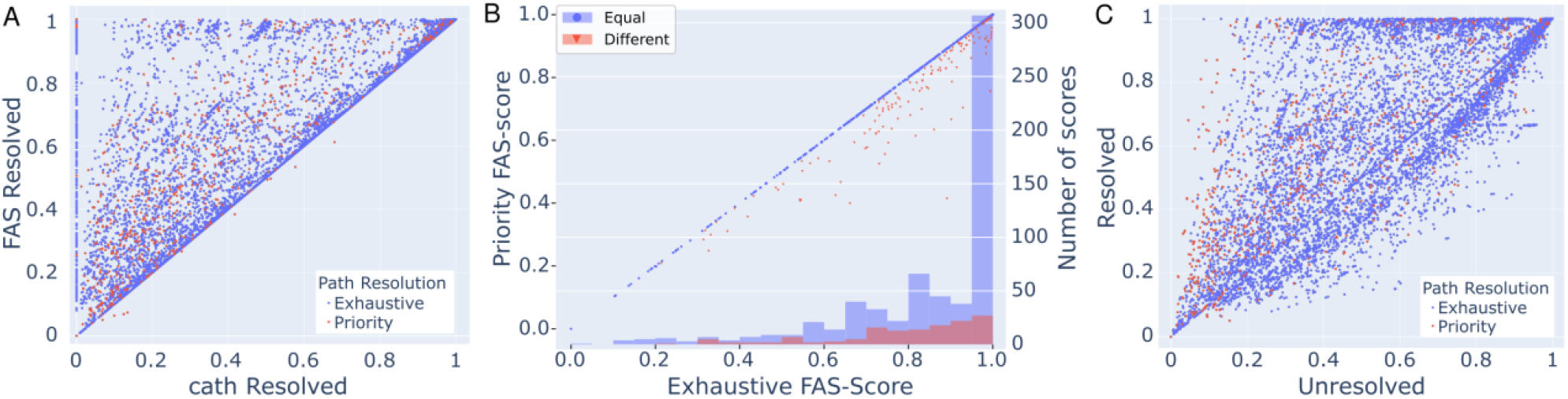
Impact of the scoring system on the architecture similarity assessment. (A) The scatter plot shows for each human-yeast ortholog pair the MLFA similarity resolving overlaps once with FAS and once with cath-resolve-hits. Architectures resolved in exhaustive mode are colored blue, those resolved with priority mode are colored red. (B) Comparison between exhaustive and priority mode. The scores are depicted in a scatterplot with the exhaustive scores on the x-axis and priority score on the y-axis (right). Protein pairs achieving similar scores with both overlap resolution methods (ΔFAS < 0.01) are represented in blue, all others in red. The histogram provides the number of proteins for both categories in 20 equally spaced FAS score bins. (C) Comparison of MLFA similarity scores using once the FAS resolved, and once the unresolved architectures (*cf*. Koestler et al. 2010). Dots above the angle bisector from 0 to 1 indicate pairs where the resolved feature architecture results in a higher FAS score, and dots below the diagonal represent the reverse case. The color code resembles that of subfigure (A).

#### 2.4.2 Priority Mode

The runtime complexity of the exhaustive search scales exponentially with the number of alternative paths in each architecture. This precludes the analysis of proteins with highly redundant architectures, of which Titin (Q8WZ42) with 10^173^ alternative paths is the most extreme example in the human protein set (Fig. S2). To cope with such complex architectures, we have devised an iterative search heuristic. We first resolve the redundant parts of a MLFA greedily by selecting at each graph junction the feature type that maximizes the partial FAS score up to and including the current feature instance. This quickly determines a lower bound for the optimal FAS score. However, it neglects that the number of instances for the selected features in the not yet visited part of the architecture graph affects both feature weighting and scoring (see Eqs. 2 and 4). To search for higher scoring alternative paths, we repeat the graph traversal this time assigning one feature type ‘priority’. At each junction, an instance of this feature type is given precedence, if it is present. Otherwise, FAS defaults to the greedy approach for this junction. We iterate over all feature type with multiple instances in the architecture and select the resolved architecture that maximizes the FAS score. The runtime complexity of the priority mode increases only linearly with the number of alternative paths (see Fig. S3). By default, FAS switches to the priority mode when the number of possible path combinations exceeds 500.

### 2.5 Output

For each evaluated pair of proteins, FAS will output their feature architecture similarity score together with the corresponding overlap-resolved feature architectures in tsv format.

## 3 Materials & Methods

### 3.1 Data

We downloaded pair-wise orthology assignments between human and yeast (*Saccharomyces cerevisiae*) created by OMA, InParanoid, and Ensembl Compara from the OrthoBench websites. We further downloaded for *Xenopus tropicalis* three QfO reference proteomes (releases 2019, 2020, 2021) from https://www.ebi.ac.uk/reference_proteomes/ as well as the proteome from NCBI RefSeq Genome (GCF_000004195.4).

### 3.2 Semantic similarity of GO annotation

For the Ensemble Compara set we downloaded for each human-yeast ortholog pair the corresponding GO term annotation (sub-ontology “Molecular Function”) (Carbon et al., 2021) from the Gene Ontology web sites (release 2022-06-22). Pair-wise semantic similarities of the GO annotations were computed with the simReL method (Schlicker et al, 2006).

### 3.3 Feature architecture differences between different gene annotations

We downloaded the full set of orthologous groups from the OMA database and selected the subset having at least nine of the following ten species represented: *Homo sapiens, Mus musculus, Rattus norvegicus, Monodelphis domestica, Gallus gallus, Pelodiscus sinensis, Anolis carolinensis, Xenopus laevis, Latimeria chalumnae, Ciona intestinalis*. Each OMA group was then extended with orthologs from the four *X. tropicalis* proteomes using fDOG (https://www.github.com/bionf/fdog) with *X. laevis* as the reference. Pair-wise FAS scores between all orthologs and the *X. laevis* reference protein were computed with FAS and the resulting data presence-absence patterns of orthologs were visualized with PhyloProfile (Tran et al., 2018).

### 3.4 Hardware

Computations were run on an Intel(R) Core(TM) i5-3470 CPU @ 3.20GHz using 1 core. FAS completed overlap resolution and similarity scoring for the non-redundant list of 14,434 ortholog pairs in 23 minutes and 22 seconds resulting in an average run-time per protein pair of less than 0.1 seconds.

## 4 Results

FAS determines the pairwise similarity between two MLFAs by resolving overlaps using a score maximization algorithm. To evaluate FAS, we used human-yeast ortholog assignments that have been submitted to the Quest for Ortholog (QfO) benchmark server (Altenhoff et al., 2020). We compared OMA (2,595 pairs; (Altenhoff et al., 2019)), InParanoid (4,578 pairs; (Sonnhammer & Östlund, 2015)), and Ensemble Compara (12,676 pairs; (Yates et al., 2016)). FAS scores were calculated for all orthologous pairs using default parameter values and a reference-based feature weighting.

### 4.1 Impact of overlap resolution on the architecture similarity score

We first investigated the effect of overlap resolution on the similarity scores using the Ensemble Compara ortholog pairs with overlaps in at least one architecture as test data (10,155 pairs). For each pair, we computed the similarity scores once resolving overlaps with the score maximization approach and once by selecting the feature instance with the lowest e-value (Lewis et al., 2019). For more than half of the investigated protein pairs (7,032), the score maximization approach resulted in higher similarity scores (ΔS_FAS_: max=1.0, median=0.09, mean=0.20, Fig. 3A). For 3,113 pairs the similarity scores were identical, and only in 10 instances the score maximization approach resulted in a lower score. Common to all ten cases was the use of the priority mode due to the high number of possible paths (Table S1). Repeating the scoring for these protein pairs enforcing the exhaustive path search revealed always scores equal or higher than the ones achieved with the e-value based overlap resolution.

We next assessed how often, and to what extent, the priority mode resulted in an underestimation of the feature architecture similarity. We computed FAS scores for 780 human-yeast protein pairs using once the exhaustive path search and once the priority mode (Fig. 3B). We selected these pairs using two criteria: (i) the proteins had at least two alternative paths through the redundant parts of their architecture, and (ii) the exhaustive path search could be completed within 1 hour on a single CPU (representing about 10^10^ path comparisons). The results from the two search modes agree in most cases (ΔS_FAS_<0.005 for 641 pairs). In 112 cases, the priority mode resulted only in a slight underestimation (0.005 < ΔS_FAS_ < 0.1) with a mean difference in the FAS score of 0.06 and a median difference of 0.04. Only for 27 protein pairs, the FAS score computed with the priority mode was off by more than 0.1. Thus, we conclude that the priority mode gives a reasonably accurate approximation of the optimal FAS score.

In the last step, we assessed the effect of resolving redundant architectures with the score maximization approach on the similarity score compared to the use of unresolved architectures (Fig. 3C). In most cases, overlap resolution resulted in higher FAS scores (5,235; ΔS_FAS_ > 0.01), but surprisingly, the opposite was true for 3,123 ortholog pairs. The priority mode was used only in 104 out of the 3,123 pairs, and thus cannot explain this observation. Taking a closer look at the corresponding architectures revealed that the FAS scores from overlapping architectures can also be artificially inflated (Fig. S4). Redundant annotations in the Pfam/SMART layer increase the total number of feature types in the feature architecture. At the same time, this results in a decrease of each individual feature weight because the sum of weights over all features in the architecture is scaled to 1 (see section 2.3.3). Therefore, the impact of a missing feature on the score is buffered by the agreeing but redundant feature types in the unresolved part of the architecture.

### 4.2 Example application 1 – FAS scoring detects differences between ortholog search tools

FAS allows the integration of feature architecture similarities into large-scale comparative sequence analysis workflows. We investigated how the choice of the ortholog search tool affects the MLFA similarity distribution between the resulting human-yeast ortholog pairs. Since the FAS score is not symmetric (see Eqs. 2 and 3), we used once the human and once the yeast sequence as the reference. Ortholog pairs identified by OMA (Altenhoff et al., 2019) have with 0.90 (0.96 with yeast as the reference) the highest median FAS score followed by those identified with InParanoid (Ref_human_: 0.80; Ref_yeast_: 0.90) (Sonnhammer & Östlund, 2015). Note the higher median FAS score for the search using the yeast protein as reference. Because FAS penalizes the absence of reference features in the target, this indicates that yeast feature architectures tend to be simpler and are often nested within more complex human architectures (see Fig. S5 for an example). Ensembl Compara orthologs have a median FAS score of only 0.62 irrespective of the reference choice (Fig. 5B). This reveals that many of the sequences that were assigned as orthologs by Ensembl Compara differ in their feature architecture to an extent that is rarely seen for orthologous pairs identified by the other two tools. Moreover, the mutually low FAS scores indicates that both the human and the yeast protein harbor features in their architecture that are not represented in the ortholog of the respective other species (see Fig. S6 for an example). The total feature counts in the human and yeast architectures for the ortholog pairs assigned by the three tools are in line with this hypothesis (Fig. 5B). In the case of OMA and InParanoid only 30.2% and 27.0% of the yeast proteins, respectively, comprise more features than their human ortholog. This number rises markedly to 39.0% for the Ensembl Compara ortholog pairs. At the same time the fraction with the same number of features is the lowest among the three tools.

**Figure 4:**
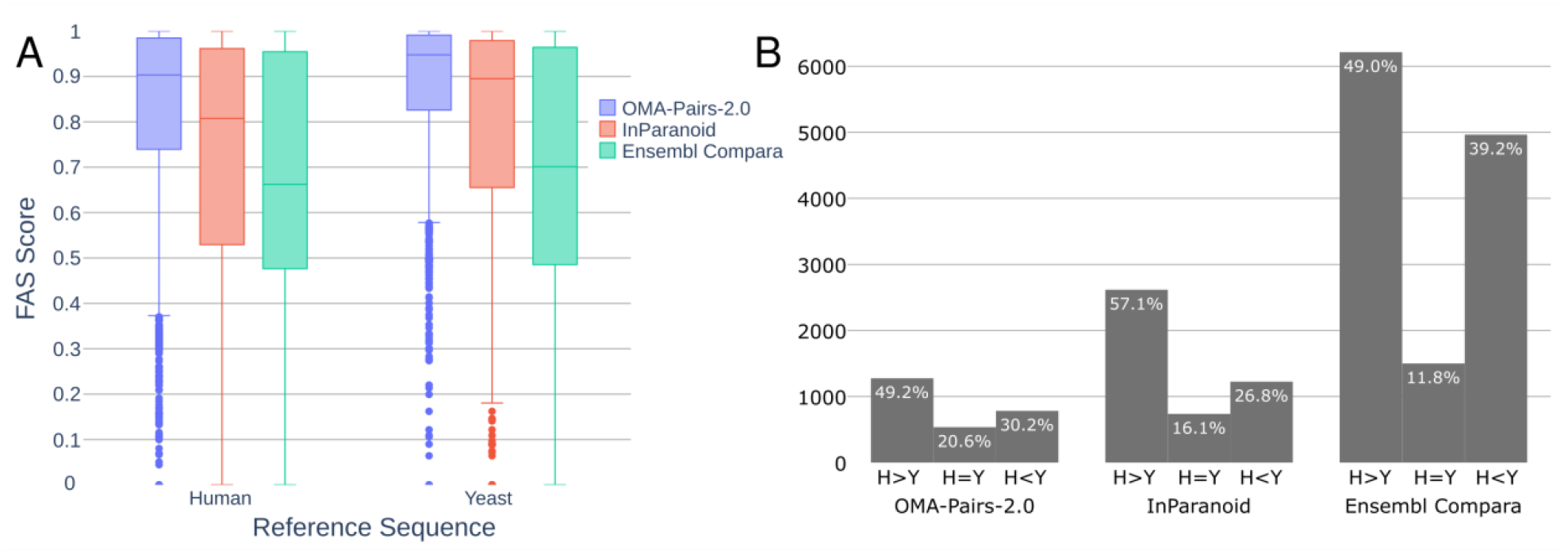
Orthology assignment tools differ in their spectrum of feature architecture similarities. The FAS score distributions for human-yeast ortholog pairs assigned by OMA (2,595 pairs), InParanoid (4,578 pairs) and Ensembl Compara (12,676 pairs) is shown in (A). (B) reveals the fraction of orthologs where the MLFA of humans comprises more features (H>Y), an equal number (H=Y) or a lower number (H<Y) than that of the yeast ortholog

**Figure 5:**
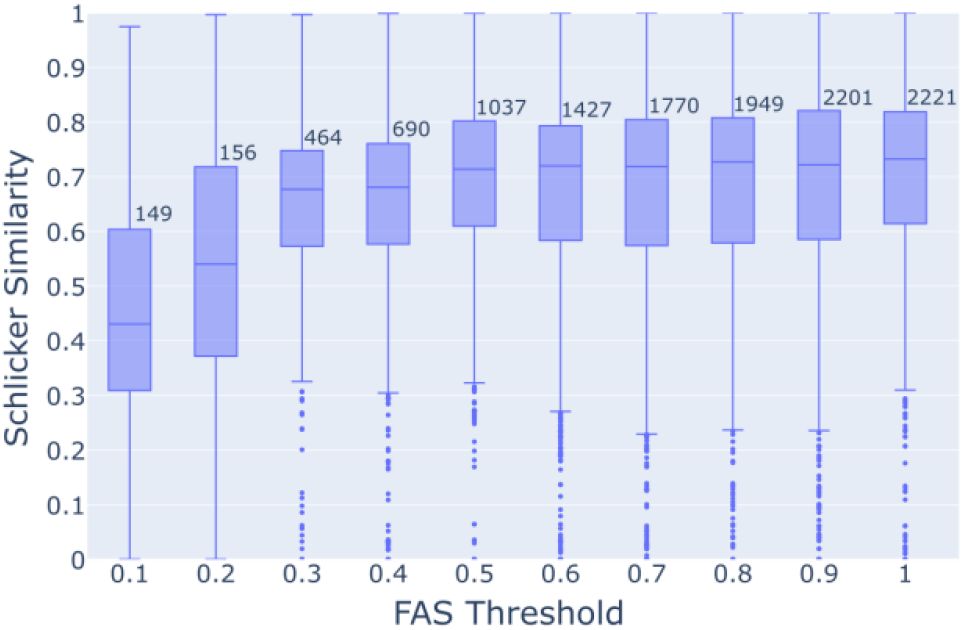
Semantic similarity of GO annotation decreases with decreasing feature architecture similarity. Ensembl Compara orthologs between human and yeast were distinguished into 10 bins based on their mean bi-directional FAS score (0 ≤ x ≤ 0.1, up to 0.9 < x ≤ 1.0). For each bin, we plotted the semantic similarities of the GO annotations (Schlicker Score) for the pairwise orthologs as a box plot. Bin sizes are indicated above each box plot

### 4.3 Low FAS scores as an indicator of functional diversification

We next investigated whether differences in the feature architecture coincide with differences in the functional description of the proteins. For each ortholog pair (Ensemble Compara), we correlated the semantic similarity of the GO term annotation (Schlicker score) with the corresponding FAS score. Distributing the ortholog pairs over ten FAS score bins in steps of 0.1 revealed that the semantic similarity of the functional annotations decreases with decreasing FAS score (Fig. 6). However, Schlicker scores vary substantially within each bin. To track down the underlying reasons, we investigated 80 extreme cases in greater detail ((|S_FAS_ – S_schlicker_| >= 0.75; Tables S2 and S3). In most cases, inaccuracies in the GO term assignment explain the deviation from a positive correlation between FAS scores and Schlicker scores (see supplementary text for two examples). However, we present two notable exceptions below.

**Figure 6:**
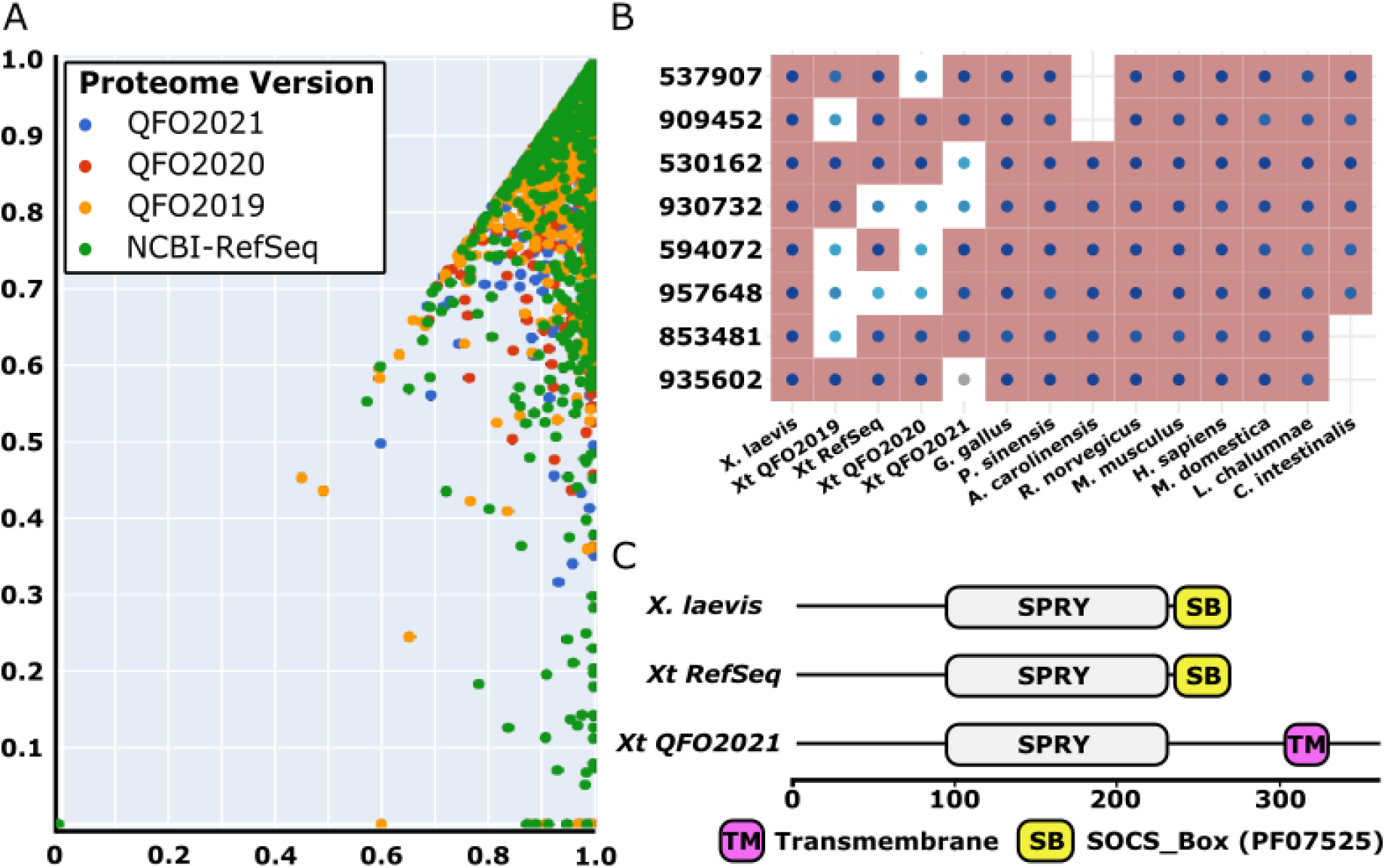
FAS scoring reveals feature architecture differences between different versions of the *Xenopus tropicalis* proteome. (A) Scatter plot of the maximum and the minimum FAS score between the reference protein of *X. laevis* and its orthologs in the four versions of the *X. tropicalis* proteome. Dot color indicates the proteome version that provides the ortholog with the lowest FAS score. (B) Section of the phylogenetic profile highlighting examples where the feature architecture differs between individual versions of the *X. tropicalis* proteome. Dots indicate the presence of an ortholog in the respective species. Dot color informs about the average bi-directional FAS score between the ortholog and the *X. laevis* reference protein ranging from 1 (dark blue) to 0.35 (grey). Cells with a white background indicate orthologs that differ significantly in their feature architecture from the reference protein. (C) Feature architecture differences in the core group 935602. The Xt_QFO2021_ ortholog (Uniprot Id: A0A6I8T262) harbors a transmembrane domain instead of a SOCS_Box Pfam domain that is present in both the Xt_RefSeq_ ortholog (Uniprot Id: XP_002936603.1) and the *X. laevis* reference (Uniprot Id A0A1L8FM06). Xt – *Xenopus tropicalis*

#### Scenario 1: High FAS - low Schlicker

Two proteins Q9Y696 (human) and Q12390 (yeast) contrast an identical feature architecture (FAS score = 0.98) with a low Schlicker score (0.01). Both proteins harbor two Pfam domains that are characteristic of glutathione-S-transferases (GST_N and GST_C), and the yeast protein is annotated as a glutathione-S-transferase (GST; EC2.5.1.18)(Ma et al., 2009). Surprisingly, the human protein is a member of the chloride intracellular channel protein family (CLIC4; Fig. S7) (Littler et al., 2005). The remarkable similarities of GSTs and CLICs on the levels of domain architecture and 3-D structure has been discussed before, e.g., (Ponsioen et al., 2009), which excludes an annotation artefact as an explanation. Instead, GSTs and CLICs are a showcase example that proteins can indeed radically differ in function without modifying their feature architecture. However, among the 80 investigated examples, this protein pair is the only of its kind. Thus, functional diversification in the context of conserved feature architectures appears rather an exception than a rule (see Supplementary Text 1 and Fig. S8).

#### Scenario 2: Low FAS – high Schlicker

Substantial differences in the feature architectures indicate a functional diversification, whereas high Schlicker scores suggest the opposite. Q7Z2Y5 (human) and Q12469 (yeast) are protein kinases catalyzing the same reaction (EC: 2.7.11.1), and are involved in cell signaling (Lin et al., 2009). Accordingly, their GO term annotation is identical. What is then the relevance of the low similarity on the feature architecture level (Fig. S6)? Both proteins share the presence of a Pkinase domain (PF00069). However, this domain is located at the N-terminus of the human protein, and at the C-terminus of the yeast protein. The human protein, which is about 1,000 amino acids longer harbors additionally a CNH domain (SMART: SM00036), which probably acts as a regulatory domain and might be involved in macromolecular interactions (Chen et al., 1999). The yeast protein harbors additionally a P21-Rho-binding domain (PF00786) that binds Rho-like small GTPases, and the N-terminus is occupied by a Pleckstrin homology domain (PH; SMART: SM00233), PH-domains are commonly found in eukaryotic signaling proteins and seem to play a role in recruiting proteins to different membranes (Wang & Shaw, 1995). These differences indicate that although both the human and the yeast protein catalyze the same reaction in the context of signal transduction, they very likely differ in their precise functions.

Cases where a low FAS score contrasts a high Schlicker score challenge the assumption that deviating feature architectures are a good proxy for the functional divergence of two orthologs. Our curation, has indicated that it is mostly poor GO annotations that drives this observation (see supplementary text; Table S3 and Fig. S9). Comparing Schlicker scores between orthologous proteins using once the full set of GO terms, and once only the GO terms with experimental validation reveals as a general trend that Schlicker scores decrease in part substantially when using experimentally verified GO terms only (Fig. S10).

### 4.4 Feature architecture similarities reveal deviating gene annotations

The extent of feature architecture change between related proteins provides an indication of their functional diversification in the courses of evolution. In the last analysis, we changed scope and demonstrate the application of FAS scoring for the detection and evaluation of differences in protein feature architectures as a consequence of alternative gene annotations. We used the African clawed frog *Xenopus tropicalis* as an example and compared four versions of its proteome, three provided as Uniprot reference proteomes (QfO2019, QfO2020, QfO2021), and one provided by the NCBI RefSeq genome database. We searched for *X. tropicalis* orthologs to 3,460 tetrapod core genes in each of the four proteome versions and evaluated their FAS score using the corresponding *Xenopus laevis* ortholog as a reference.

Figure 6A shows for each gene the minimal and the maximal FAS score obtained with the orthologs from the four alternative proteomes. This revealed in part substantial differences in the feature architectures between the different versions of the *X. tropicalis* proteins. Overall, the RefSeq version of the *X. tropicalis* proteome provided annotations that are most consistent in their feature architecture with the orthologs in other species (Table 1). However, the QfO2021 version is only marginally worse, whereas the QfO2020 version of the *X. tropicalis* proteome had the most proteins with deviating architectures. Within the QfO reference proteomes, we note a general trend of domain architectures becoming more consistent with the *X. laevis* reference protein with newer releases (Fig. 6B; Table 1). However, there are individual examples for the opposite case (Figs. 6B and C). This indicates that MLFA comparisons can be used to rapidly identify even rare cases where the annotated gene structure should be revised.

**Table 1:** FAS scores between the reference protein of *X. laevis* and its orthologs in the four versions of the *X. tropicalis* proteome.

| Proteome | Orthologs | FAS<sub>Max</sub> | FAS<sub>>0.1</sub><sup>\$</sup> | FAS<sub>Δ0.1</sub><sup>#</sup> | FAS<sub>Min</sub><sup>§</sup> |
| --- | --- | --- | --- | --- | --- |
| NCBI_RefSeq | 3132 | 1829 (58%) | 9 | 234 (7.5%) | 208 (6.6%) |
| QfO2021 | 3089 | 1359 (43%) | 5 | 169 (5.5%) | 121 (3.9%) |
| QfO2020 | 2600 | 1161 (44%) | 6 | 183 (7.0%) | 145 (5.6%) |
| QFO2019 | 3073 | 1237 (40%) | 19 | 232 (7.5%) | 182 (5.9%) |

<sup>\$</sup> Number of orthologs where FAS<sub>Max</sub> exceeds the runner up from a different proteome version by at least 0.1 score units
<sup>#</sup> Number of orthologs with a FAS score of at least 0.1 score units below the maximal score achieved with an ortholog from a different proteome version
<sup>§</sup> Number of orthologs achieving the smallest FAS score across the four proteome versions that is at least 0.1 scoring units below FAS<sub>Max</sub>

## 5 Discussion

Protein feature architectures (FAs) provide relevant information when assessing whether two evolutionarily related proteins have functionally diverged or may still share the same ancestral function. The gold standard in interpreting architecture similarities is still cost- and labor-intense human curation. The scoring scheme developed here aims at reducing the gap between an automatically generated pairwise FA similarity assessment and their visual interpretation of FA changes facilitating the use of FAs also in the analysis of large cohorts of orthologs.

The main innovation in FAS is the score maximization algorithm to resolve overlapping feature types. We could show that this allows to identify orthologs with deviating feature architectures more reliably than existing scoring schemes using either an e-value based naïve resolution (Lewis et al., 2019), or the use of unresolved architectures (Koestler et al., 2010). Overlaps within the Pfam/SMART layer in the MLFA are generated by more than one pHMM with a significant similarity to the same amino acid sequence. Within Pfam, such pHMMs are often grouped in clans (Finn et al., 2006). Thus, the re-coding of pHMMs in an architecture with their clan assignment can ameliorate, at least partly, the overlap resolution problem. However, the functional spectrum of clan members can be substantial. For example, the Zinc beta-ribbon clan (CL0167; 89 members) harbors 17 and 21 pHMMs representing different kinds of zinc-ribbon and zinc finger domains, respectively. This set of short, nucleic acid binding domains is complemented among others with eight pHMMs representing different ribosomal proteins and one model of a large-conductance mechanosensitive channel. Treating these functionally diverse models in an architecture comparison as synonymous to zinc-ribbon or zinc finger domains unduly reduces the sensitivity to detect architecture differences. Moreover, clans are restricted to pHMMs provided by Pfam, and overlaps between redundant pHMMs provided by different databases cannot be resolved.

This latter problem can, in principle, be addressed by using the Conserved Domain Database (CDD) as an annotation resource (Lu et al., 2020). Protein domain superfamily provided by CDD result from a manually curated clustering of protein domain models that annotate overlapping footprints on protein sequences. Version 3.19 of this database comprises 62,852 models from a broad collection of source databases, *e.g*., Pfam v.32, SMART v6.0, COGs v1.0, TIGRFAMs v15 and Entrez Protein Clusters) and collapses them into 4,617 multi-model superfamilies. The integration of CDD superfamilies as an additional feature class into FAS is straightforward. However, here we decided to demonstrate the integration of individual source databases on the fly. As one main advantage, FAS can thereby always use the latest release of the individual source databases, and to complement this information with custom models trained by the individual users.

To demonstrate the use of the FAS, we have shown that the distribution of MLFA similarities between pairs of human-yeast orthologs depends on the tools that have been used for the orthology assignments. Orthology inference is an evolutionary reconstruction problem, and thus a ground truth does not exist. To still assess performance differences between individual orthology assignment tools, the Quest for Orthologs benchmark service currently provides 12 challenges, comprising various discordance tests between gene trees reflecting the evolutionary histories of the predicted orthologs and the corresponding expected tree topologies, the agreement in Enzyme Commission (EC) numbers, and the semantic similarity in GO term annotation (Altenhoff et al., 2020). Following the idea that orthologs tend to be functionally more conserved than paralogs (Tatusov et al., 1997), the latter two challenges correlate the accuracy of the orthology assignments for two proteins with their functional similarity. However, only a fraction of proteins are enzymes, and using the semantic similarity of GO term annotations as a proxy for functional equivalence bears many pitfalls (Thomas et al., 2012; our own results). Moreover, orthologs can functionally diverge (see Fig. S7). MLFA similarities reflected in the FAS score distribution may therefore constitute a relevant complementary challenge in the ortholog benchmark that is more comprehensive than EC classification and less susceptible to annotation biases than GO term assignments.

The tracing of functionally equivalent orthologs across taxa is essential for a reliable protein annotation transfer (*e.g*., Aramaki et al., 2020; Cantalapiedra et al., 2021; Kanehisa et al., 2016). In turn, the identification of functionally diverged orthologs can help unravelling evolutionary changes that account for lineage-specific phenotypic characteristics, e.g., that differentiate pathogens from their non-pathogenic relatives. We could show that low FAS scores readily identify ortholog pairs with a strong indication for a functional diversification. However, errors during gene annotation such as the missing of individual exons, or the artificial fusion or fission of genes generate the same signal. While it is common practice to use protein sets from related organisms to guide the identification of genes in a newly sequenced organism (*e.g*., Braker2 pipeline (Brůna et al., 2021)), testing the resulting gene predictions for consistency across taxa is not. Existing approaches focus mainly on the length of the resulting proteins but not on their feature architecture (Seppey et al., 2019). Comparing orthologs from four versions of the *X. tropicalis* proteome and up to 10 further representatives spanning the vertebrate diversity revealed proteome-specific deviations in otherwise evolutionarily highly conserved feature architectures. These instances represent in most cases artefacts in gene annotation, and unless they become corrected, provide a spurious signal of lineage-specific functional diversification. With the help of FAS, it is straightforward to identify and subsequently correct such errors in the annotation of protein coding genes.

### 5.1 Limits

The similarity assessment of proteins using their feature architectures still bears a prominent limitation: the information gained from different feature architectures varies depending on the architecture complexity. The impact of a gain or a loss of the same feature type on the FAS score decreases with increasing complexity of the feature architecture. A case-by-case adjustment of feature weights can ameliorate this effect if the feature type, e.g., a transmembrane domain, is known. However, in large-scale screens prevents the varying granularity of the FAS score the *a priori* specification of a score threshold below which functional diversification has to be assumed. Here, manual curation of feature architectures optimally in the context of FAS score-aware phylogenetic profiles (see Fig. 6; Birikmen et al. 2021) is still necessary.

## Acknowledgements

This work was supported by the Research Funding Program Landes-Offensive zur Entwicklung Wissenschaftlich-ökonomischer Exzellenz (LOEWE) of the State of Hessen, Research Center for Translational Biodiversity Genomics (TBG).

