## Supplemental text, Supplemental Figs. S1 - S10, Supplemental Tables S2 - S2 for "FAS: Assessing the similarity between proteins using multi-layered feature architectures"

### Supplementary material

#### Supplementary text

A high feature architecture similarity is typically considered as evidence that two evolutionarily related proteins are at least similar in their function. In turn, pronounced differences in the feature architectures should indicate a functional diversification. We evaluated to what extent the FAS score that reflects the pairwise similarity between two multi-layered feature architectures and the Schlicker score that reflects the semantic similarity of the functional annotation with GO terms that have been assigned to the two proteins are correlated. While we see a strong trend that proteins with a low FAS score tend to have a low Schlicker score (see Fig. 5 in main text) there is a surprisingly high variation within the individual FAS bins. We evaluated 80 examples with an extreme difference between the two scores. In most cases, limitations in the GO annotation can explain the difference (see Table S2 and main text), and we discuss two examples in greater extent.

Scenario 1: High FAS – low Schlicker scores. Agreeing feature architectures are often taken as evidence that two orthologs share the same or at least a similar function. Q5TAP6 (human; gene name: UTP14C) and Q04500 (yeast; gene name: UTP14) contrast a high FAS score (0.90) with a Schlicker score of only 0.06. The feature architecture of the yeast protein differs only in the presence of an N-terminal D rich extension from that of its human ortholog. In line with their conserved architectures, both proteins are annotated as a U3 small nucleolar RNA-associated protein 14, and both are components of the small subunit processome (Black et al., 2018) (human - <https://www.genenames.org/data/gene-symbol-report/#!/hgnc_id/HGNC:20321>). Thus, there is no indication that the two proteins have diverged in function. Instead, the low Schlicker score reflects a poor GO annotation of the two proteins (Table S2) rather than a functional diversification.

Scenario 2: Low FAS – high Schlicker scores. Substantial differences in the feature architectures indicate a functional diversification, whereas high Schlicker scores suggest the opposite. Three yeast proteins, Q06328, P38279 and Q12010, are annotated as amino acid transmembrane transporters. Accordingly, their FAs harbor several transmembrane domains (Fig. S9, Table S2). The human ortholog A1A4F0 is less than half the size of the yeast proteins and its FA is devoid of any features, which results in a FAS score of 0. Despite the lack of transmembrane domains, A1A4F0 is annotated with GO terms associated with amino acid transmembrane transport (Evidence code IBA: inferred from biological aspect of ancestor; ECO:0000318). This explains the high semantic similarities of the GO annotations for these protein pairs (0.97), which must be considered as spurious according to our evidence.

#### Supplementary Figures

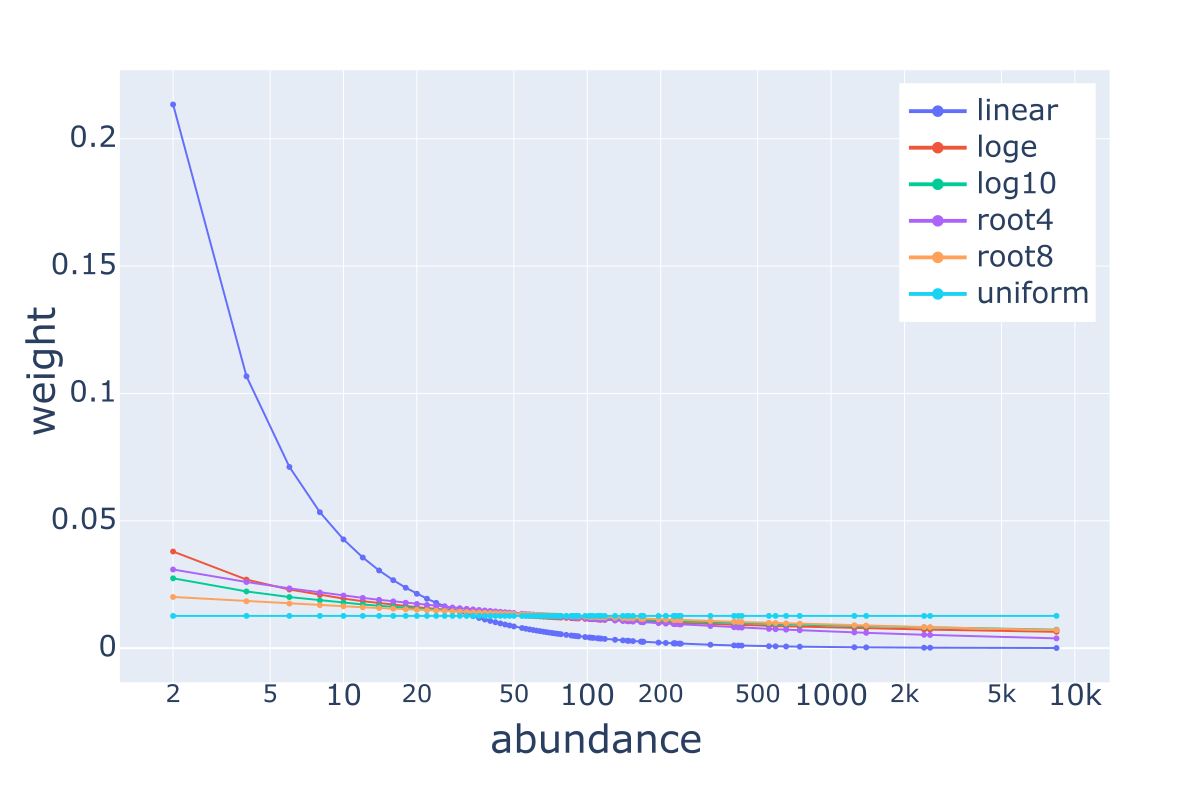

Figure S1: Impact of the weighting function on the feature weights in dependency of the feature abundance. Six different weighting functions are currently implemented in FAS (see plot inlay). Linear weighting strongly biases the weighting in favor of rare features, whereas uniform weighting renders the feature weight independent of the feature abundance. ‘Abundance’ gives the number of proteins in the yeast proteome carrying at least one instance of a feature type.

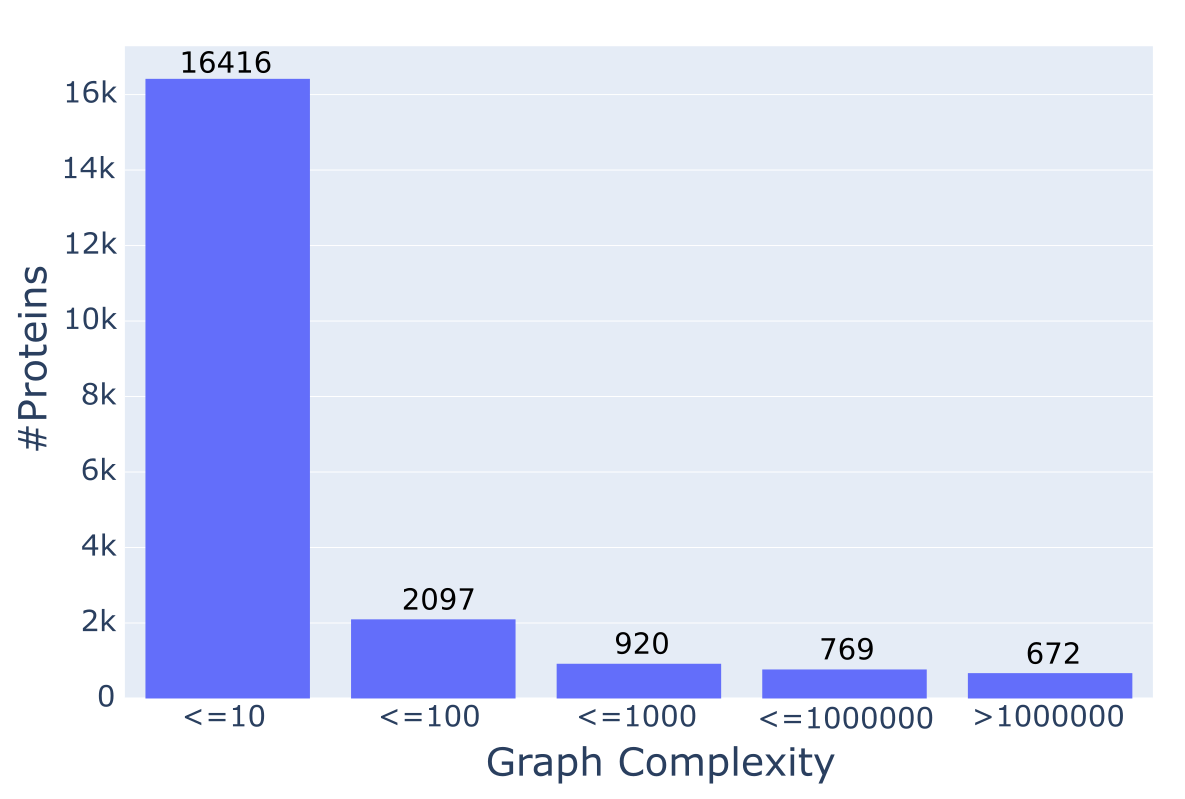

Figure S2: Distribution of graph complexities across the human protein set (QfO 2020 Reference Proteome). We evaluated the number of alternative paths through redundant parts of an MLFA and refer to this as graph complexity. The histogram shows that the vast majority of proteins has 10 or fewer alternative paths. However, still 672 proteins have more than 1 million alternative paths.

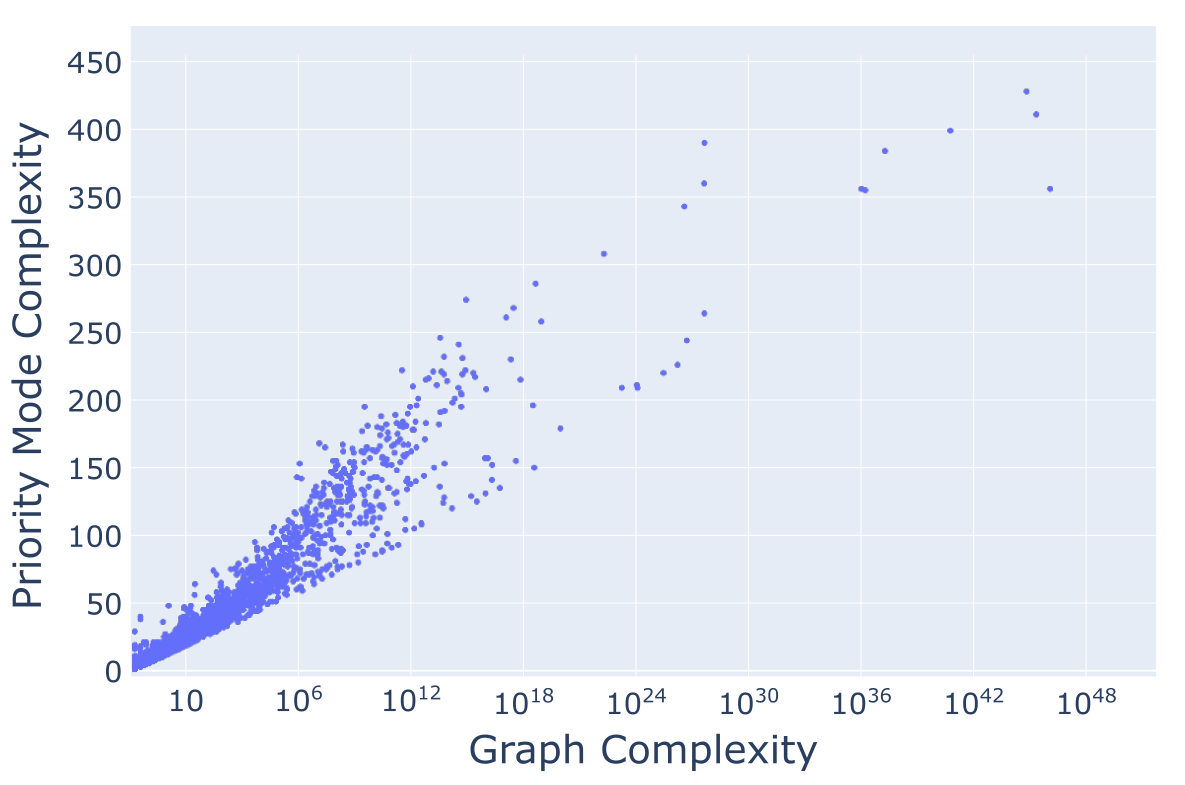

Figure S3: Path search complexity of the exhaustive vs. the priority mode. Each dot represents a human protein. The x-axis gives the number of possible representative paths through overlapping parts of the feature architecture (Pfam-/SMART layer). The y-axis gives, for the same protein, the number of paths that have to be evaluated when using the priority mode for finding the path that maximizes the FAS score.

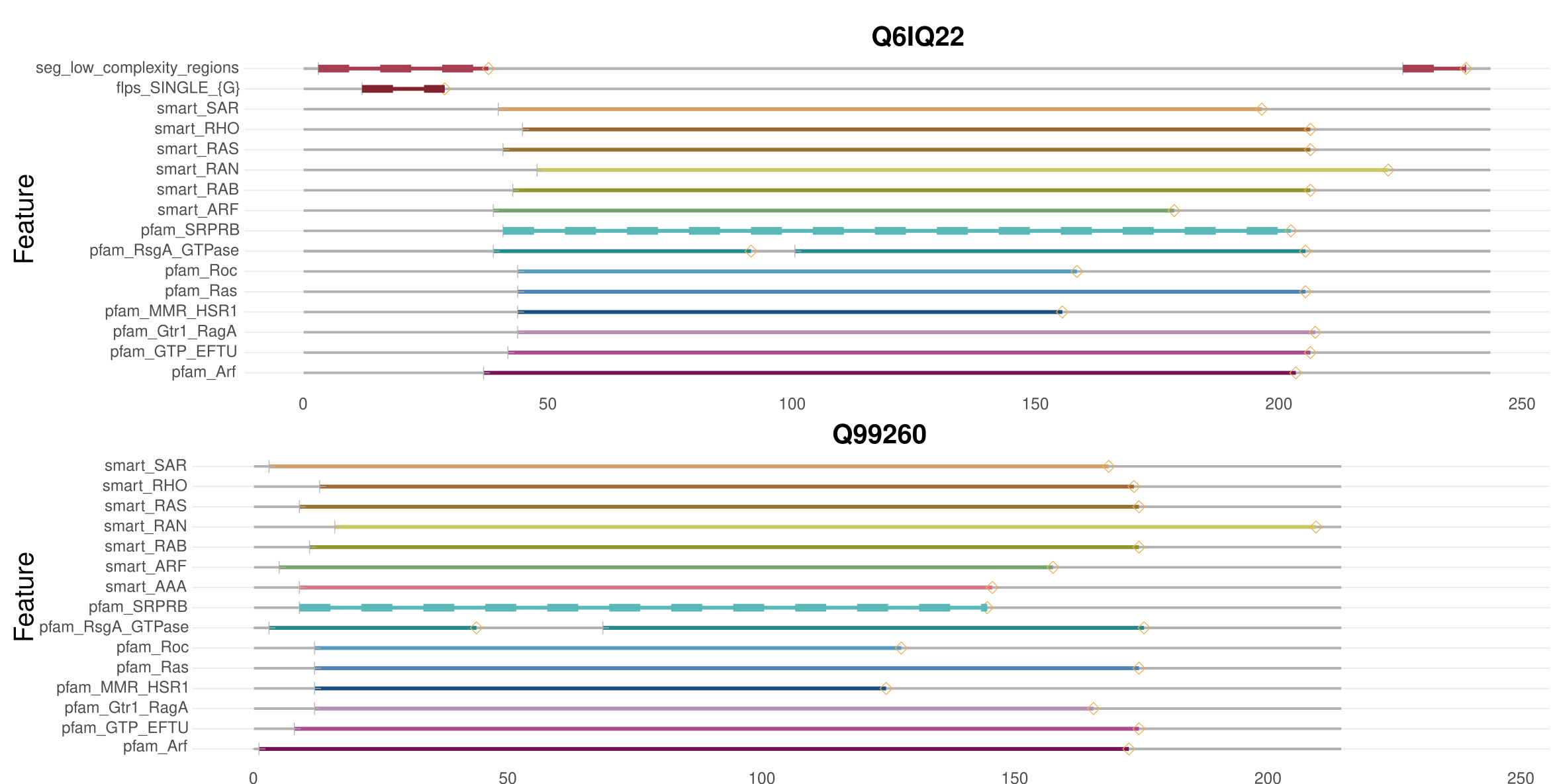

Figure S4: Feature architectures of a protein pair for which the similarity score is higher when using the unresolved architectures. The central part of both proteins is congruently annotated with a multitude of overlapping Pfam- and SMART domains that, when remain unresolved, contribute to more than 90% of the similarity score. Consequently, the differential presence/absence pattern of low complexity regions in the N- and C-terminal parts of the two proteins reduces the similarity score only to a minor extent. Resolving the overlaps in the Pfam-/SMART layer removes this buffering effect, and thus the similarity score decreases as the architecture differences due to the low complexity regions present only in the human protein (Q6IQ22) increase in their weight

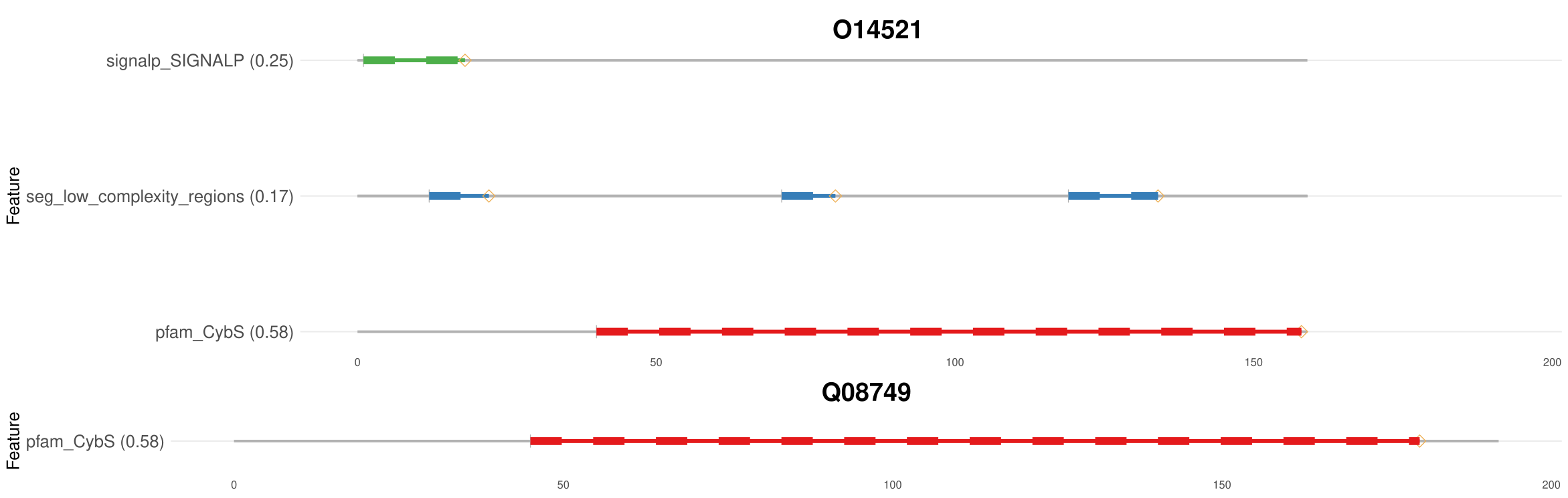

Figure S5: An example of nested feature architectures. The yeast protein (Uniprot-ID: Q08749) harbors only an Pfam CybS domain. The human protein (Uniprot-ID: O14521) carries additionally an N-terminal signal peptide as well as three low complexity regions. This difference results in a lower FAS score when using the human protein as the reference, while it is close to 1 when using the yeast protein as the reference

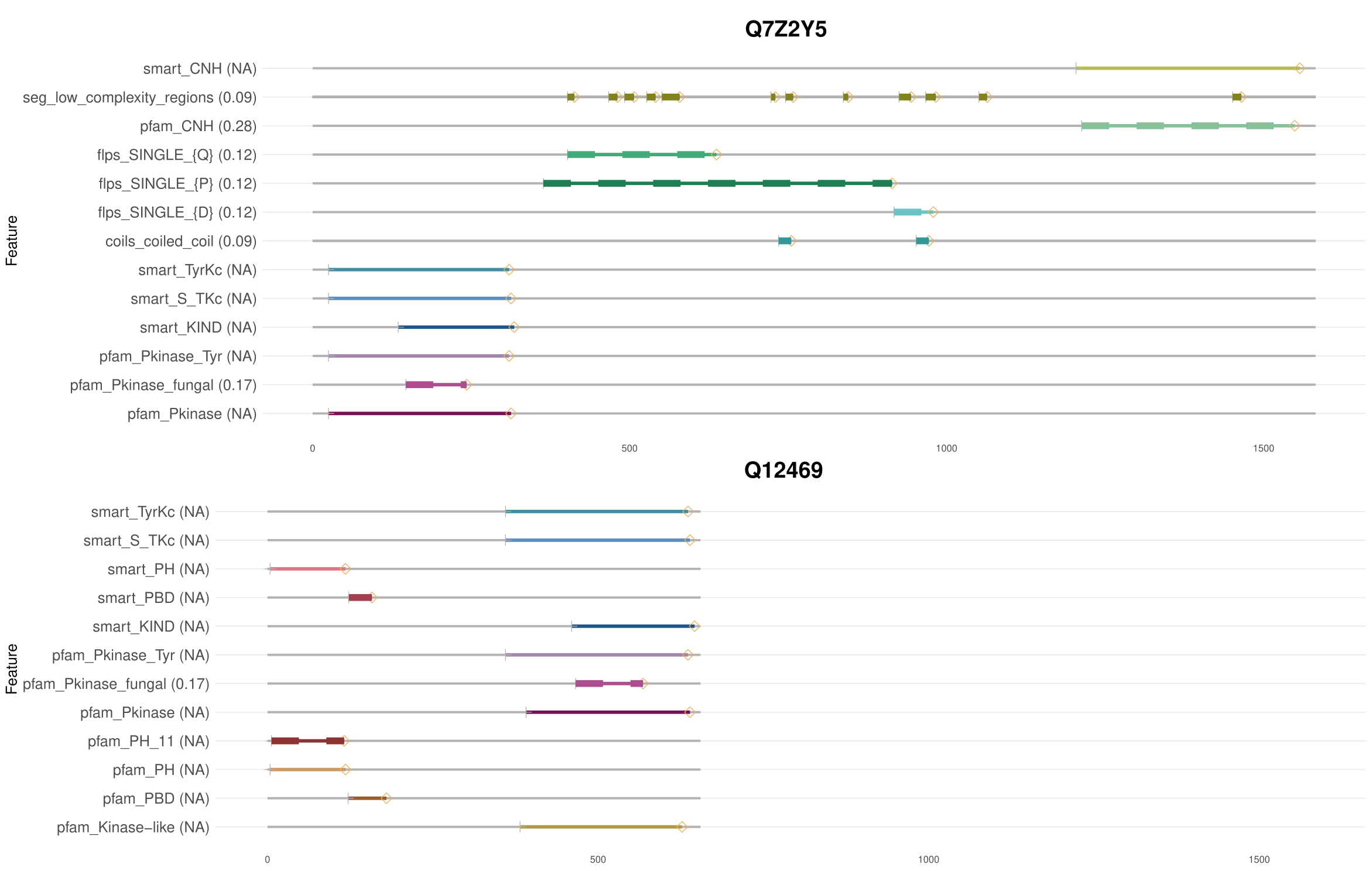
Figure S6: Feature architectures of Q7Z2Y5 and Q12469. Both proteins have a similar GO annotation (Schlicker Score: 0.97) but differ substantially in their feature architecture. On the feature architecture level, both proteins share only the presence of a tyrosine kinase domain, and correspondingly the FAS score is only 0.2

A

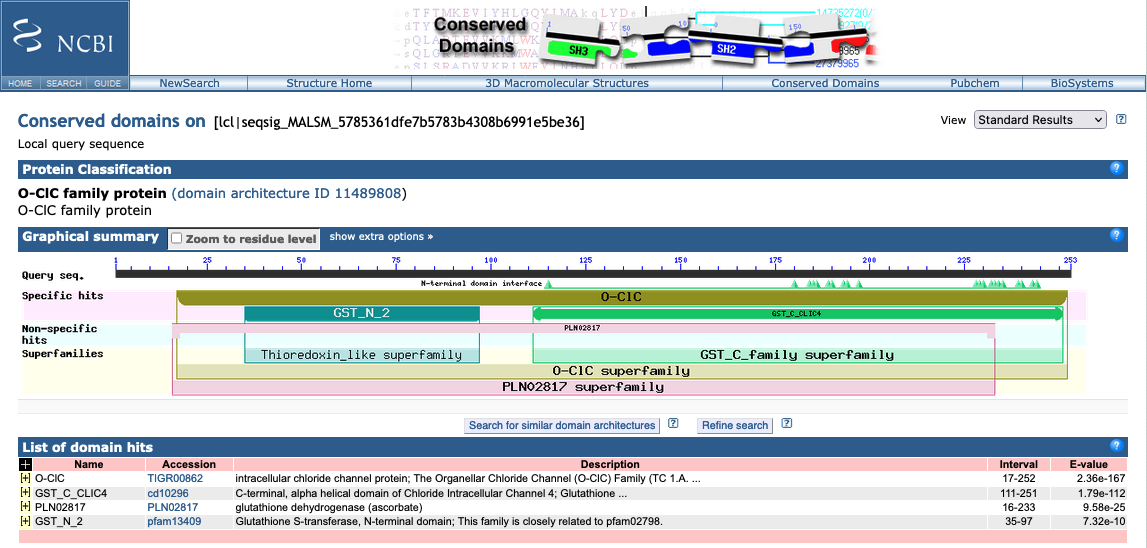

A
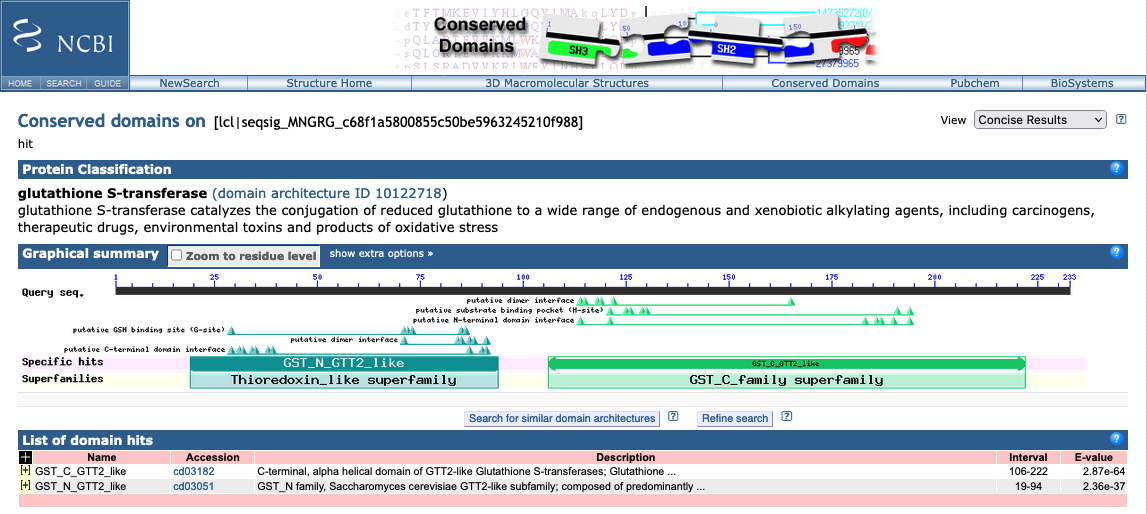
Figure S7: CD search result for Q9Y696 and Q12390. The human protein Q9Y696 is annotated as a chloride channel protein (A), the yeast protein Q12390 is annotated as a gluthatione-S-transferase (B)

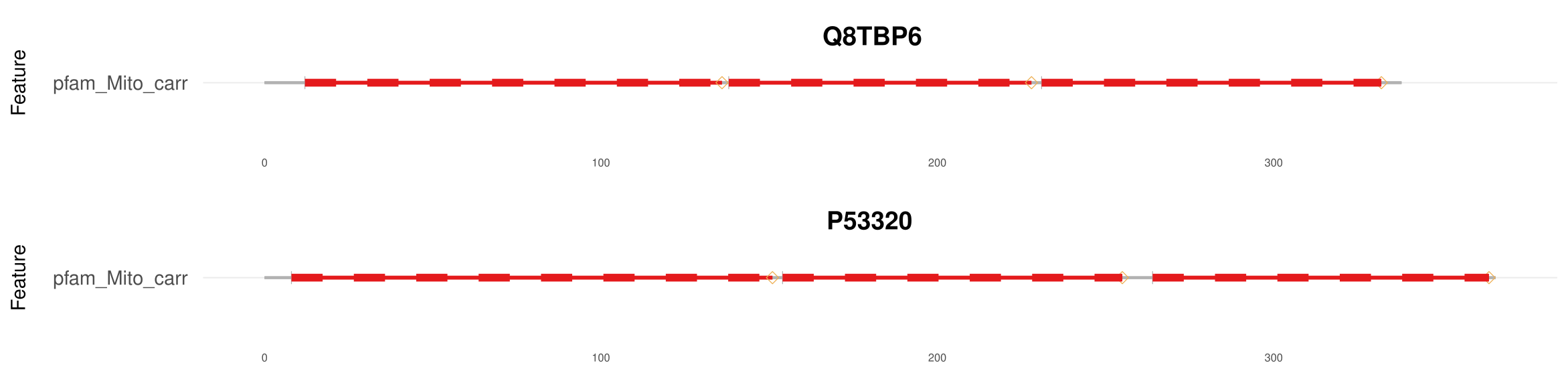
Figure S8: Feature architectures of Q8TBP6 and P53320. The two proteins act as mitochondrial transporters both in human and yeast and share identical feature architectures. Still, the Schlicker score of the corresponding GO annotations is 0.0 (Table S2)

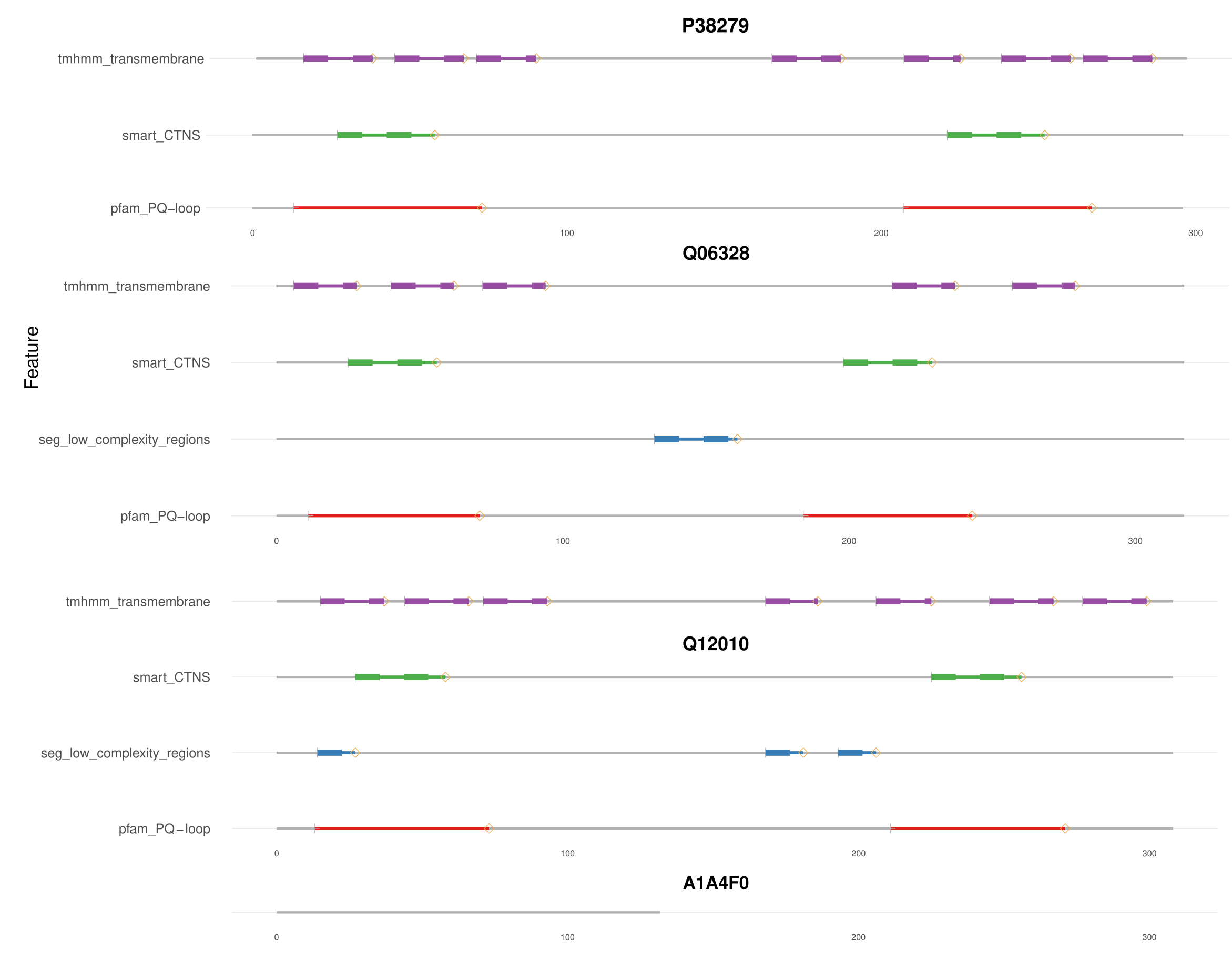
Figure S9: Examples for proteins that share the same GO annotation despite pronounced differences in their feature architectures. Three yeast proteins (Q12010, Q06328 and P38279) display almost identical feature architectures. Their human ortholog (A1A4F0) is devoid of any features, and hence its FAS score when compared to the yeast proteins is 0. The discrepancy in feature architecture between the human and yeast proteins is contrasted by a Schlicker score of 0.97 suggesting that all four proteins share the same function (Table S2; see Supplementary Text)

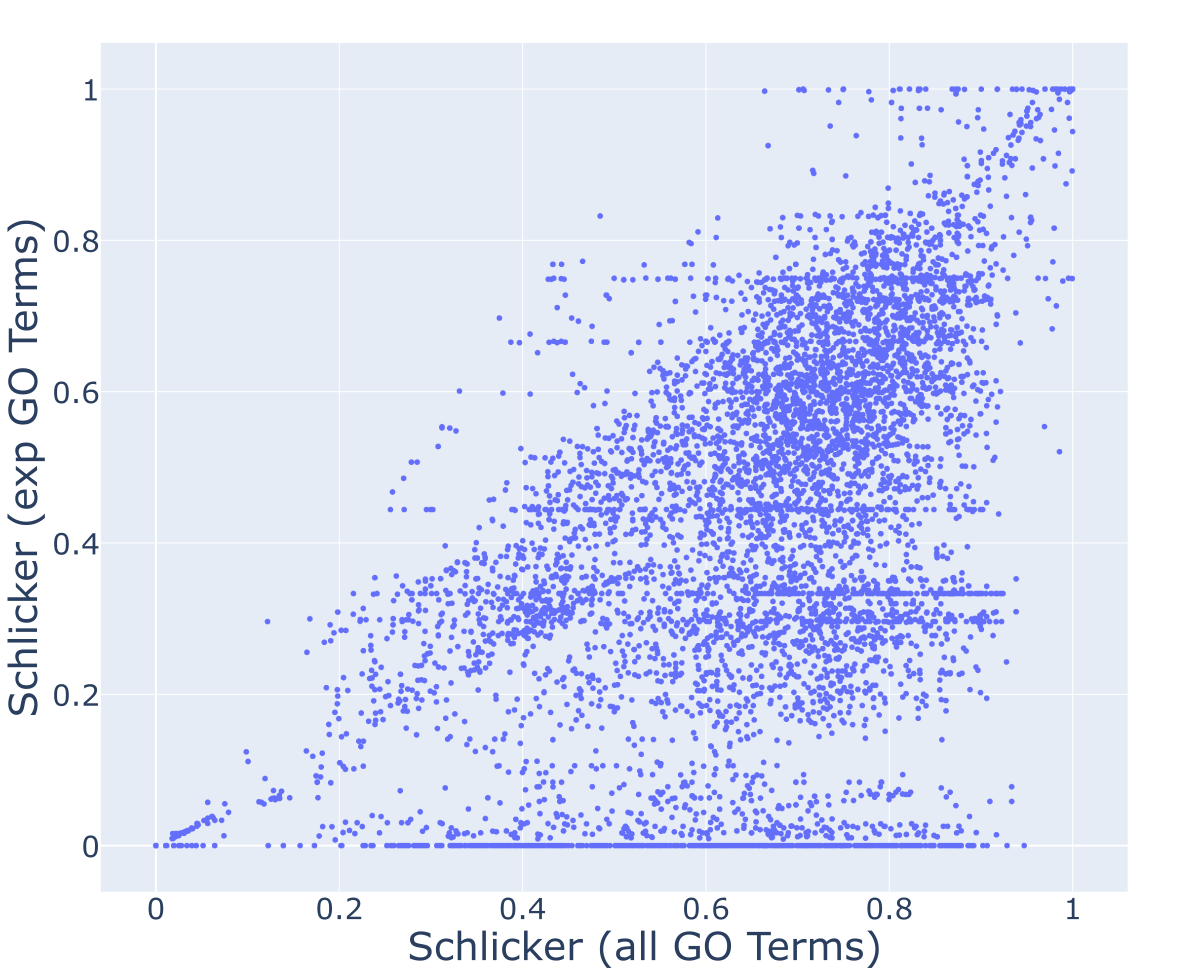

Figure S10: Semantic similarities of GO annotations using once all GO terms and once only GO terms with experimental evidence. The scatterplot shows the semantic similarities (Schlicker score) of Ensembl Compara ortholog pairs decreases when only GO terms with experimental evidences are used

#### Supplementary tables

##### Table S1: Path complexities of the 10 human-yeast ortholog pairs for which the score maximization approach to resolve overlaps resulted in a lower score than the e-value-based approach

| Human ID  (Uniprot) | Yeast ID  (Uniprot) | # of alternative paths (human) | # of alternative paths (yeast) | # of path evaluations (exhaustive) |
| --- | --- | --- | --- | --- |
| Q12913 | P25044 | 9216 | 6 | 55296 |
| Q14554 | Q12404 | 1703 | 6 | 10218 |
| Q8IWX7 | Q12118 | 216 | 291 | 62856 |
| Q92824 | P09232 | 630686 | 1 | 630686 |
| Q92824 | P25036 | 630686 | 1 | 630686 |
| Q92824 | P25381 | 630686 | 1 | 630686 |
| Q96EQ0 | P15705 | 391 | 5893568 | 2304385088 |
| Q96EQ0 | P38825 | 391 | 222870 | 87142170 |
| Q9HD43 | P25044 | 15008 | 6 | 90048 |
| Q9Y4ES | P39933 | 4096 | 236520 | 968785920 |

##### Table S2: Molecular Function GO term annotation for the orthologs (Q8TBP6 & P53320), (A1A4F0 & Q06328, P38279, Q12010) and (Q5TAP6 & Q04500)

| Protein | GO Term ID | Description |
| --- | --- | --- |
| Q8TBP6 | GO:0005347 | ATP transmembrane transporter activity |
| P53320 | GO:0030170 | pyridoxal phosphate binding |
| A1A4F0 | GO:0015174 | basic amino acid transmembrane transporter activity |
| A1A4F0 | GO:0015189 | L-lysine transmembrane transporter activity |
| Q06328 | GO:0015174 | basic amino acid transmembrane transporter activity |
| P38279 | GO:0015174 | basic amino acid transmembrane transporter activity |
| Q12010 | GO:0015174 | basic amino acid transmembrane transporter activity |
| Q5TAP6 | GO:0005515 | protein binding |
| Q04500 | GO:0003674 | molecular function |
| Q04500 | GO:0005524 | ATP binding |

##### Table S3: Evaluation of 80 human-yeast ortholog pairs with discrepant FAS and Schlicker scores (|S_FAS_ – S_Schlicker_| >= 0.75)

See separate file
